# Born to be young: prenatal thyroid hormones increase early-life telomere length

**DOI:** 10.1101/2020.05.07.083667

**Authors:** Antoine Stier, Bin-Yan Hsu, Coline Marciau, Blandine Doligez, Lars Gustafsson, Pierre Bize, Suvi Ruuskanen

**Affiliations:** Department of Biology, University of Turku, Turku, Finland; Institute of Biodiversity, Animal Health and Comparative Medicine, University of Glasgow, Glasgow, UK; Department of Biometry and Evolutionary Biology, CNRS, Université de Lyon France; Department of Ecology and Genetics / Animal Ecology, University of Uppsala, Uppsala, Sweden; School of Biological Sciences, University of Aberdeen, Aberdeen, UK

**Keywords:** Aging, mitochondria, telomere length, bird, fetal programming

## Abstract

Prenatal environmental conditions can have lifelong consequences on health and aging. The underlying mechanisms remain nonetheless little understood. Thyroid hormones (THs) are important regulators of embryogenesis transferred from the mother to the embryo. In an avian model, we manipulated embryo exposure to maternal THs through egg injection and investigated the consequences on postnatal growth and aging. We first report that mitochondrial DNA (mtDNA) copy number and telomere length significantly decrease from early-life to late adulthood, thus confirming that these two molecular markers are hallmarks of aging in our wild bird model. The experimental elevation of prenatal THs levels had a transient positive effect on postnatal growth. Elevated prenatal THs had no effect on mtDNA copy number but significantly increased telomere length both soon after birth and at the end of the growth period (equivalent to offsetting *ca.* 4 years of post-growth telomere shortening). These findings suggest that prenatal THs have a key role in setting the ‘biological’ age at birth, and thus might influence longevity.

---

Prenatal environmental conditions can have lifelong consequences on health and aging, but most remains to be done to uncover the mechanisms linking the pre and postnatal stages (1). Thyroid hormones (THs) are master regulators of development, health and aging (2–4). They are transferred from the mother to the embryo (4) and thyroid disorders during pregnancy can induce developmental pathologies in human (5). Surprisingly, there is currently no experimental data on the effects of prenatal THs on postnatal health and aging.

Two challenges must be overcome when testing for long-term consequences of maternally-transmitted hormones. First, manipulating the prenatal hormonal environment in mammalian models is usually problematic since treatments are applied to the mother and can have indirect effects on the embryo. Avian models offer an ideal alternative since prenatal conditions can be directly manipulated through hormonal injection in the egg (4). The second challenge is to be able to measure long-term effects using health and aging markers that are accurately and adequately mirroring age-related health impairments. Two promising markers are mitochondrial DNA (mtDNA) copy number and telomere length since both markers are decreasing with age and have been associated with increased mortality risks (6, 7). Telomere length is considered as a proxy of ‘biological’ age, with more relevance to mortality risk than chronological age (8). Interestingly, most of the inter-individual variation in telomere length is already set at birth, and thus may be caused by different exposure to maternal hormones (9).

In a free-living bird population of collared flycatchers *(Ficedula albicollis),* we first investigated age-related changes in telomere length and mtDNA copy number using data covering the entire lifespan spectrum for this population. Results show that mtDNA copy number (Fig. 1A) and telomere length measured either using a relative qPCR method *(rTL,* Fig. 1B) or an absolute *in-gel* quantification *(absTL,* Fig. 1C) significantly decreased from growth completion to late-adulthood. These findings are in accordance with reports from the human literature (6, 7) and validate their use as hallmarks of aging in our avian model.

**Fig. 1:**
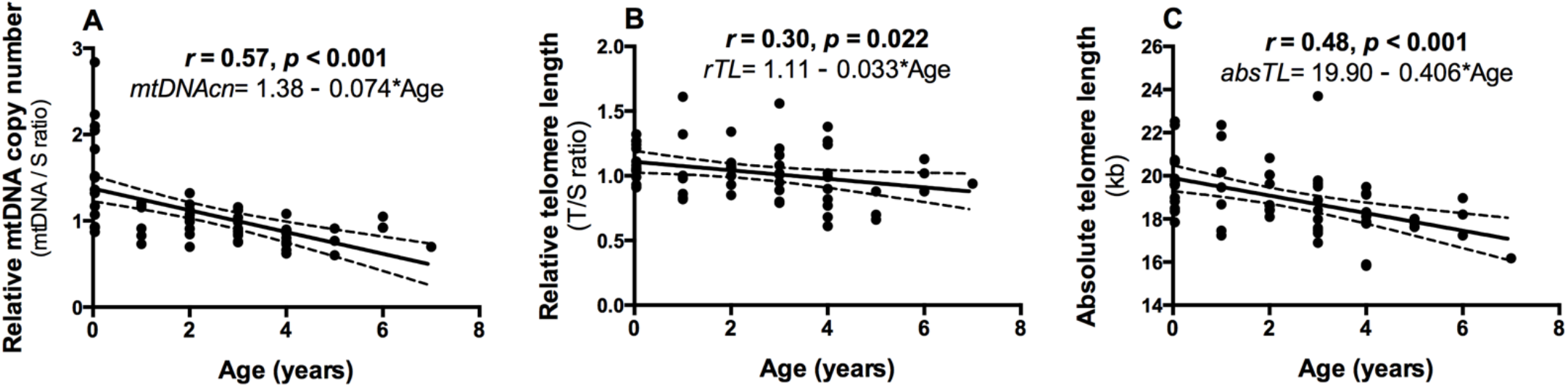
Age-related variation in potential key hallmarks of aging in wild collared flycatchers: **(A)** decrease in relative **mtDNA copy number**, **(B)** decrease in **relative telomere length** measured with qPCR, and **(C)** decrease in **absolute telomere length** measured with *in-gel* TRF. Data is cross-sectional, adult birds were of known-age and chicks were 12 days old (from control group only, 1 chick per nest). Regression lines are plotted ± 95% C.I., N = 58 (44 adults, 14 nestlings).

Using egg-injection of THs we then investigated the effect of prenatal THs on postnatal mtDNA copy number and telomere length. Based on the known stimulation of mitochondrial biogenesis by THs (10), we predicted that increasing prenatal THs should increase early-life mtDNA copy number, which could be a cellular pathway supporting the transient growth-enhancing effect previously demonstrated (11). Conversely, since THs are known to increase oxidative stress and enhance growth (two pathways accelerating telomere shortening (9)), we predicted that increasing prenatal TH levels should shorten telomere length at birth, and/or increase early postnatal telomere shortening.

We confirmed the growth-enhancing effect of prenatal THs (Fig. 2A) in our subsample of individuals from (11) but found no significant impact of prenatal THs on mtDNA copy number (Fig. 2B), despite a considerable early-life reduction in mtDNA copy number during the growth period (equivalent to the reduction occurring over 3.5 years in individuals post-growth, based on Fig. 1A). Contrary to our predictions, increasing prenatal THs led to longer telomeres (measured as *rTL)* soon after hatching (Fig. 2C), and this effect was maintained at the end of the growth period (day 12). This is confirmed by the analysis of absolute telomere length *(absTL)* at day 12, showing longer telomeres in birds hatched from TH-injected eggs (Fig. 2D). The effect of increasing prenatal THs on telomere length was substantial, being equivalent to offsetting *ca.* 4.3 years *(rTL)* and 3.6 years *(absTL)* of telomere shortening (based on Fig. 1B and 1C).

**Fig. 2:**
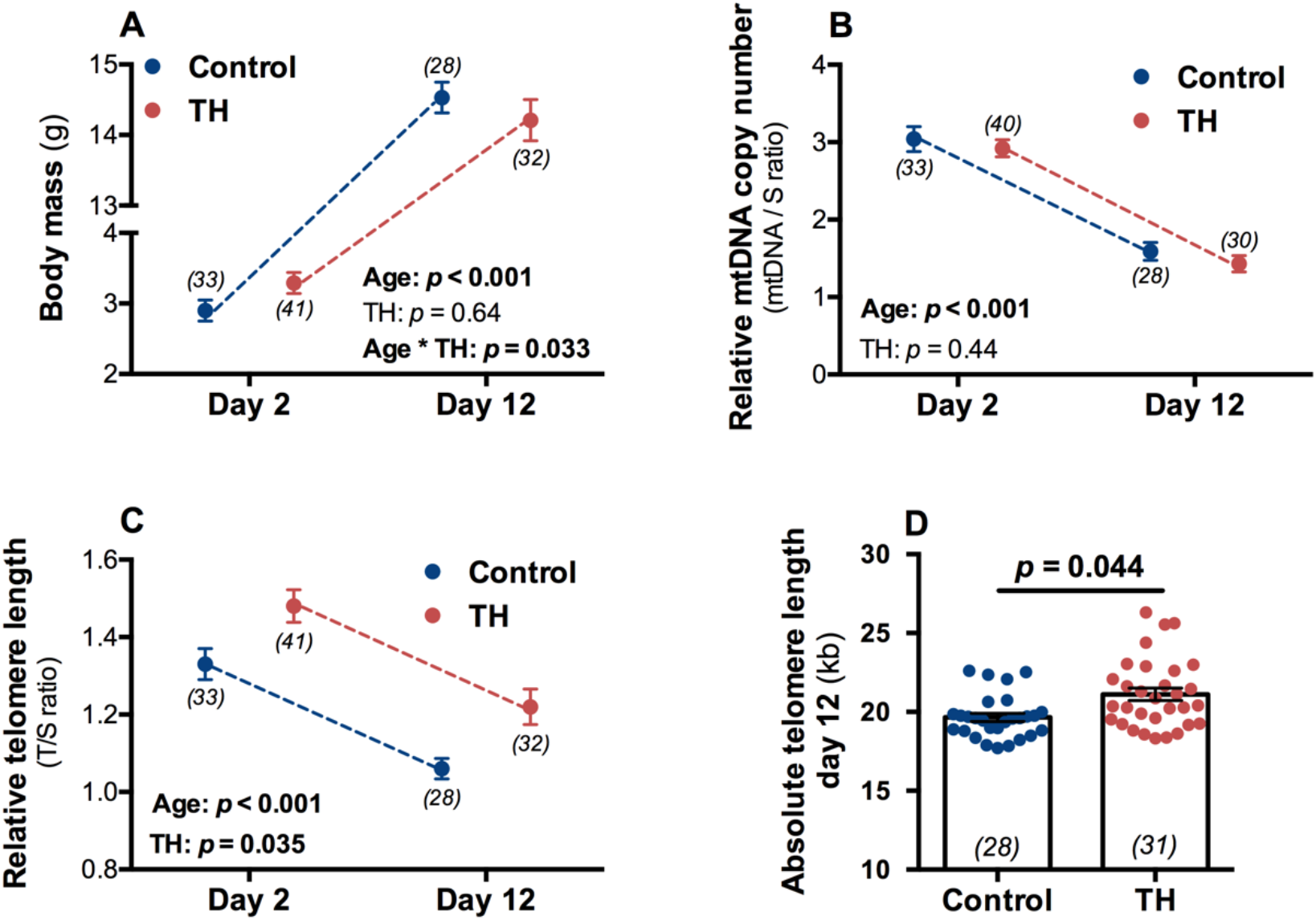
Effects of experimental prenatal thyroid hormone elevation on: **(A) body mass** growth, **(B)** early-life dynamics of **mtDNA copy number**, **(C)** early-life dynamics of **relative telomere length**, and **(D) absolute telomere length** at the end of growth (day 12). Means are plotted ± SE, *p*-values and sample sizes are indicated within each panel.

The beneficial effect of prenatal THs on telomere length is unlikely related to oxidative stress prevention, since we previously found no differences in oxidative stress markers in these experimental birds (11). One previous study reported that the promoter of *hTERT* (the catalytic subunit of the enzyme telomerase, responsible for elongating telomeres) contains a binding site for THs (12). Consequently, one hypothesis would be that prenatal THs could elongate telomeres early in life through the activation of the telomerase enzyme. The positive correlation found between the mRNA expression of the thyroid-stimulating hormone and telomere length in human adipose tissue could support such an hypothesis (13).

While the exact mechanisms remain to be identified, our study demonstrates that prenatal TH levels have the potential to elongate telomeres in early-life, and thus to set the ‘biological’ age at birth. This is the first study to show that telomere length at birth could be increased by modulating the prenatal hormonal environment. Thyroid function is known to influence cardiovascular disease risk and life expectancy in adult humans (3), but no information is currently available regarding the impact of prenatal TH exposure on adult health and lifespan. Epidemiological and long-term experimental studies investigating the impact of prenatal THs on lifespan are now required to establish if the effect observed here on telomeres translates into a longevity gain.

## Experimental procedures

The study was conducted in the long-term monitored population of collared flycatchers on Gotland, Sweden (Jordbruksverkets permit no. ID 872). We selected 44 adult birds of known-age (1 to 7 years old, *i.e.* cross-sectional data; maximum lifespan = 9.8 years) from the long-term monitoring program. Thirty-two nests were used for the prenatal manipulation of THs, 16 *Control* (vehicle-injected) and 16 *TH* nests in which eggs were injected with *ca.* a 2SD increase of TH egg content based on natural range, following the procedure described in (11). Birds were weighed and blood sampled as soon as possible after hatching (day 2, < 10μL of blood) and at the end of growth (*i.e.* day 12; < 50μL of blood). Relative mtDNA copy number of blood cells has been measured as described in (14). Both relative telomere length (*rTL* measured using qPCR) and absolute telomere length *(absTL,* measured using *in-gel* TRF) have been measured as described in (15). Age-related variations in mtDNA copy number and telomere length were tested using parametric correlation tests. The effects of prenatal TH elevation and age on body mass, mtDNA copy number and telomere length (*rTL* and *absTL*) were tested using linear mixed models, with nest identity as a random effect (to control for multiple birds per nest), bird ID as the repeated effect, and age, treatment and their interaction as fixed effects. Non-significant interactions were removed from final models. Sex was excluded from final analyses since it was never significant. Data used in this article is publicly available at: https://figshare.com/s/be8dca3133cc1db8af90.

## Acknowledgements

We are grateful to many students and Szymek Drobniak for their help in the field, as well as to Pat Monaghan for providing access to TRF facilities. The project was funded by a Marie Sklodowska-Curie Postdoctoral Fellowship (#658085) and a ‘Turku Collegium for Science and Medicine’ Fellowship to AS, and an Academy of Finland grant (# 286278) to SR.

## Notes

**Competing interest statement:** the authors declare having no competing interests

https://figshare.com/s/be8dca3133cc1db8af90

### Competing Interest Statement

The authors have declared no competing interest.

